# CD11b Activation Reduces Myeloid Brain Infiltration and Mitigates Synucleinopathy in a Model of Parkinson’s Disease

**DOI:** 10.1101/2025.08.11.669712

**Authors:** Ameera M. Shaw, Viviana Jimenez, Billy Nguyen, Jayda Duvernay, Solji Choi, Mike Cameron, Bryan A. Killinger, Vineet Gupta

**Affiliations:** Department of Neurology, Rush University Medical Center, Chicago, IL, USA 60612; Department of Molecular Medicine, Wertheim UF Scripps Institute, Jupiter, FL 33458; Department of Internal Medicine, University of Texas Medical Branch, Galveston, TX 77555

**Keywords:** neuroinflammation, CD11b, LA1, Parkinson’s disease

## Abstract

The pathology of Parkinson’s disease is defined by α-synuclein (α-syn) aggregation into neuronal Lewy bodies, which may lead to chronic neuroinflammation and dopaminergic neurodegeneration. Misfolded α-syn activates Toll-like receptor signaling in microglia, leading to downstream activation of NF-κB and subsequent release of pro-inflammatory cytokines. These cytokines recruit pro-inflammatory myeloid cells from circulation, thereby amplifying neuroinflammation. Thus, reducing microglial activation and myeloid cell infiltration has the potential to reduce neuroinflammation and PD pathology. Here, we investigated a targeted immunomodulatory strategy using LA1, a novel, small-molecule agonist of CD11b, a β2 integrin receptor highly and selectively expressed on myeloid cells and microglia. CD11b has key roles in cell adhesion, migration, and phagocytosis. Previous work has demonstrated that CD11b agonism via LA1 transiently enhances integrin-mediated adhesion that limits immune cell transmigration and tissue infiltration. CD11b agonism also suppresses TLR-driven inflammatory signaling and myeloid cell activation. To evaluate its efficacy *in vivo*, we utilized pre-clinical Parkinson’s disease model by stereotaxically delivering AAV2-SYN to induce α-synuclein overexpression in the murine midbrain. Mice were treated with oral LA1 for four or eight weeks and analyzed. LA1 treatment significantly reduced microglial activation and decreased brain infiltration of peripheral immune cells, thereby attenuating α-synuclein-induced neuroinflammation. These findings suggest that CD11b agonism may offer a dual-action therapeutic approach in Parkinson’s disease by dampening pro-inflammatory responses by central and peripheral myeloid cells.

---

Parkinson’s disease (PD) is a chronic, progressive neurodegenerative disorder affecting more than 10 million individuals globally (1). It is pathologically characterized by the selective degeneration of dopaminergic neurons in the substantia nigra pars compacta (SNpc) and the widespread accumulation of intracellular α-synuclein (α-syn) aggregates, known as Lewy pathology (2,3,4). Although the precise mechanisms underlying PD remain incompletely understood, an expanding body of literature implicates neuroinflammation as a key driver of disease progression rather than a mere byproduct of neurodegeneration (5,6).

Microglia, the innate immune cells of the central nervous system (CNS), rapidly respond to pathological stimuli such as misfolded α-synuclein, mitochondrial dysfunction, and reactive oxygen species (7). Upon activation, microglia adopt a pro-inflammatory phenotype marked by elevated expression of cytokines (e.g., TNF-α, IL-1β), reactive oxygen species, and antigen-presenting molecules such as MHCII (8,9). While microglial activation may initially serve a protective function, persistent or dysregulated activation exacerbates neuronal injury, thereby promoting a self-perpetuating cycle of neuroinflammation and neurodegeneration (10).

In addition to microglial activation, disruption of the blood-brain barrier in PD facilitates the infiltration of peripheral immune cells, particularly CCR2+ monocytes (11,12). These monocyte-derived cells can enter the inflamed brain in response to chemokine gradients (e.g., CCL2) and amplify the neuroinflammatory cascade, further contributing to tissue damage (11). While monocyte infiltration is often associated with pathogenic inflammation, there is increasing evidence that these cells exhibit plasticity and can adopt regulatory or reparative roles depending on context and signaling cues (13).

One critical mediator of myeloid cell function is the integrin CD11b (also known as αM integrin) that is selectively expressed on microglia, monocytes, macrophages and other myeloid cells. On the cell surface, CD11b heterodimerizes with CD18 to form integrin CD11b/CD18 (also known as αMβ2, CR3 and Mac-1). This integrin receptor is involved in processes including cell adhesion, migration, phagocytosis, and immunomodulation (14). Although CD11b is often associated with proinflammatory activity, its signaling is complex and context-dependent. Notably, pharmacological agonism of CD11b dampens inflammation by stabilizing the integrin in a high-affinity conformation, promoting adhesion and reducing migration and cytokine release (15,16).

Leukadherin-1 (LA1) is a small-molecule agonist that selectively targets CD11b and has demonstrated efficacy in modulating myeloid responses across multiple disease models, including experimental autoimmune encephalomyelitis (EAE), sepsis, systemic lupus erythematosus (SLE), and cancer (17,18,19,20,21). LA1 treatment has been shown to reduce peripheral immune cell infiltration, suppress pro-inflammatory signaling, and enhance phagocytic clearance functions in myeloid populations (17,18,19,20,21). These findings suggest that targeting CD11b could represent a novel immunomodulatory strategy for neurodegenerative diseases characterized by maladaptive inflammation.

In the present study, we investigated the therapeutic potential of CD11b activation in a well-established mouse model of α-synucleinopathy. Using adeno-associated virus (AAV)–mediated overexpression of human α-synuclein in the substantia nigra, we hypothesized that CD11b agonism via LA1 would attenuate both resident microglial activation and the recruitment of inflammatory peripheral myeloid cells to the CNS. This study aims to elucidate the role of CD11b signaling in PD pathogenesis and evaluate the translational potential of immune-modifying therapies in α-synuclein-driven neuroinflammation.

## RESULTS

### Orally delivered LA1 has low CNS penetration that is improved when co-administered with an efflux inhibitor

Pharmacokinetic analysis revealed that orally administered LA1 exhibited a plasma half-life of about an hour, with peak plasma concentrations occurring between thirty- and sixty-minutes post-administration (Table S1; Fig. S1). Following a single oral dose, LA1 concentrations in the kidney closely paralleled plasma levels, peaking within the same thirty-to-sixty-minute window (Fig. 1B). In contrast, LA1 levels in the brain were substantially lower, representing only 3–10% of concurrent plasma concentrations (Fig. 1B). Despite this low brain penetration, LA1 persisted longer in the brain relative to plasma, declining at a slower rate. Pretreatment with the P-glycoprotein (P-gp) inhibitor elacridar significantly enhanced central nervous system penetration of LA1, resulting in brain concentrations that more closely matched plasma levels, ranging from 60–98% of plasma concentrations (Fig. 1D). Importantly, chronic oral administration of LA1, either alone or in combination with elacridar, twice daily for eight weeks did not lead to drug accumulation in the plasma or brain, as no residual buildup was observed (Table S2).

**FIGURE 1.**
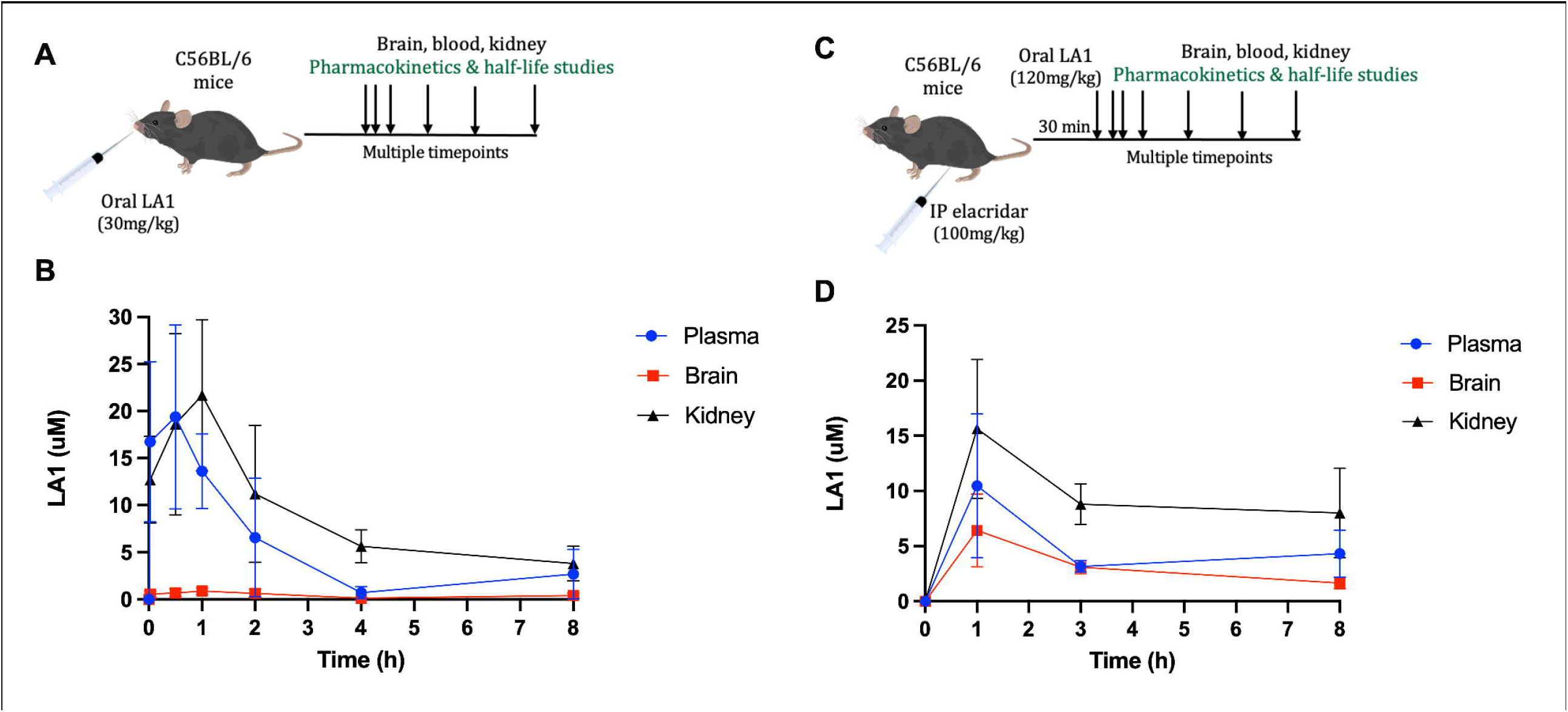
LA1 concentration in plasma and tissues over time. (A) Experimental paradigm for pharmacokinetic studies of a single oral LA1 dose. (B) LA1 concentration in plasma and tissues over time after a single oral dose. (C) Experimental paradigm for pharmacokinetic studies of a single oral LA1 dose after elacridar pretreatment. (D) LA1 concentration in plasma and tissues over time after elacridar pretreatment.

### pSer129 expression decreased with CD11b agonism

Immunohistochemical analysis of pSer129 revealed marked expression in the ipsilateral midbrain of AAV-SYN injected mice treated with vehicle for four weeks (Fig. 2A, C). Treatment with the CD11b agonist LA1 alone or in combination with the efflux inhibitor elacridar significantly reduced pSer129 expression to compared to vehicle-treated controls with a concomitant loss of nuclear hyperphosphorylation (Fig. 12A, C). Notably, nuclear pSer129 accumulation was absent in the ipsilateral striatum of AAV-SYN injected mice across all treatment groups at this time point (Fig. 2B, D). Control mice injected with PBS or AAV-GFP exhibited minimal pSer129 expression in the ipsilateral midbrain and striatum, with no evidence of nuclear hyperphosphorylation (Fig. 12). At the eight-week time point, AAV-SYN mice receiving vehicle displayed sustained nuclear hyperphosphorylation and elevated pSer129 levels in the midbrain (Fig. 3A, C). In contrast, mice treated with LA1 or LA1 + elacridar exhibited significantly reduced pSer129 expression relative to vehicle controls, along with diminished nuclear localization (Fig. 3A, C). Consistent with earlier findings, no nuclear hyperphosphorylation was observed in the ipsilateral striatum across all groups at eight weeks (Fig. 3B, D). PBS- and AAV-GFP–injected controls continued to show minimal pSer129 expression in both the midbrain and striatum at this later time point (Fig. 3).

**FIGURE 2.**
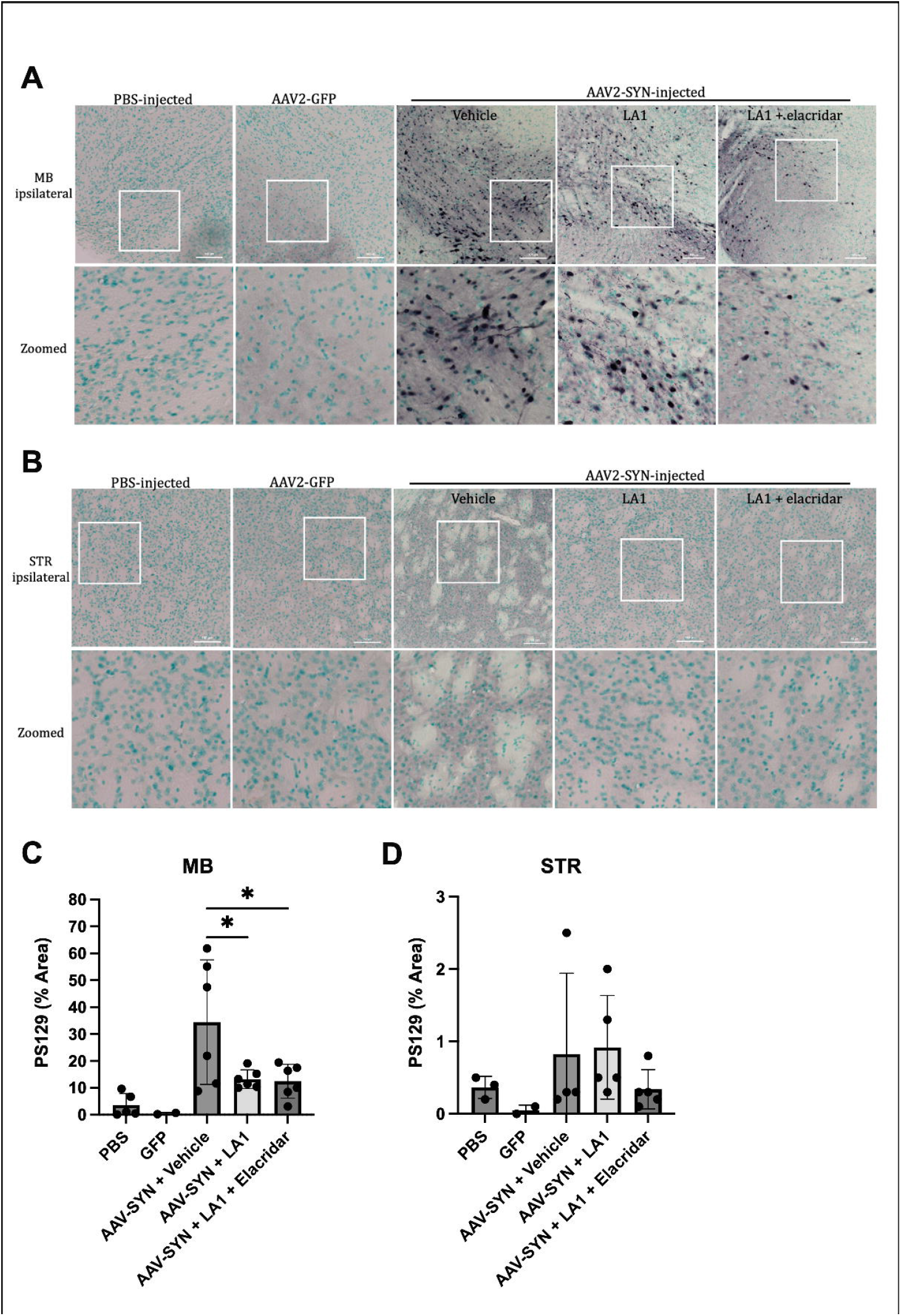
pSer129 expression decreases in the midbrain after four weeks of treatment. Representative IHC images of pSer129 expression in the ipsilateral MB (A) and STR (B) of PBS, AAV-GFP, or AAV-SYN injected mice treated with vehicle, LA1, or LA1 with elacridar for four weeks. Images taken at 20X magnification (top) and digitally zoomed (bottom). Scale bar = 100μm. Subsequent quantification of PSER129 immunoreactivity in the MB (C) and STR (D). Mean ± SEM is shown. PBS n = 5, AAV-GFP n = 2, AAV-SYN n = 6 per treatment group. *p<0.05. MB = midbrain, STR = striatum.

**FIGURE 3.**
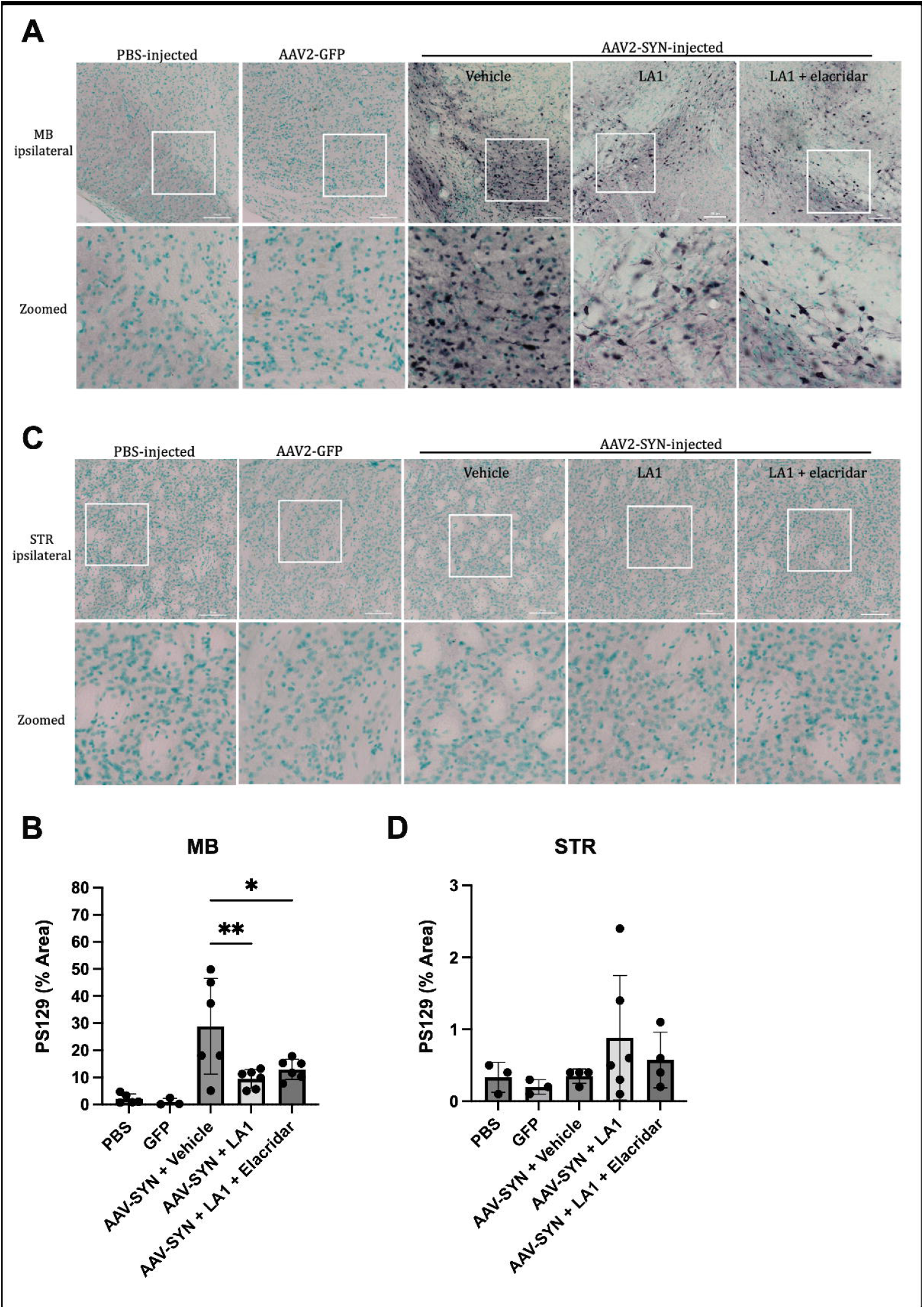
pSer129 expression decreases in the midbrain after eight weeks of treatment. Representative IHC images of pSer129 expression in the ipsilateral MB (A) and STR (B) of PBS, AAV-GFP, or AAV-SYN injected mice treated with vehicle, LA1, or LA1 with elacridar for eight weeks. Images taken at 20X magnification (top) and digitally zoomed (bottom). Scale bar = 100μm. Subsequent quantification of PSER129 immunoreactivity in the MB (C) and STR (D). Mean ± SEM is shown. PBS n = 5, AAV-GFP n = 3, AAV-SYN n = 6 per treatment group. *p<0.05, *p<0.01. MB = midbrain, STR = striatum.

### LA1 treatment decreases microglial activation and Iba1 expression

To further investigate the efficacy of CD11b agonism in attenuation α-synuclein mediated neuroinflammation, assessment of microglial activation by Iba-1 immunostaining revealed robust microgliosis in the midbrain of AAV-SYN–injected mice treated with vehicle for four weeks, characterized by increased Iba-1 expression and a bushy morphology with enlarged cell bodies and shortened, thickened processes (Fig. 4A, C). In contrast, LA1 treatment significantly reduced microglial activation, as evidenced by decreased Iba-1–positive area and attenuation of reactive morphology (Fig. 4A, C). Co-treatment with LA1 and elacridar further suppressed microglial activation, reducing Iba-1 expression to levels comparable to PBS- and AAV-GFP–injected controls (Fig. 4A, C). No significant changes in Iba-1 expression or microglial morphology were observed in the striatum across any treatment groups at the four-week time point (Fig. 4B, D).

**FIGURE 4.**
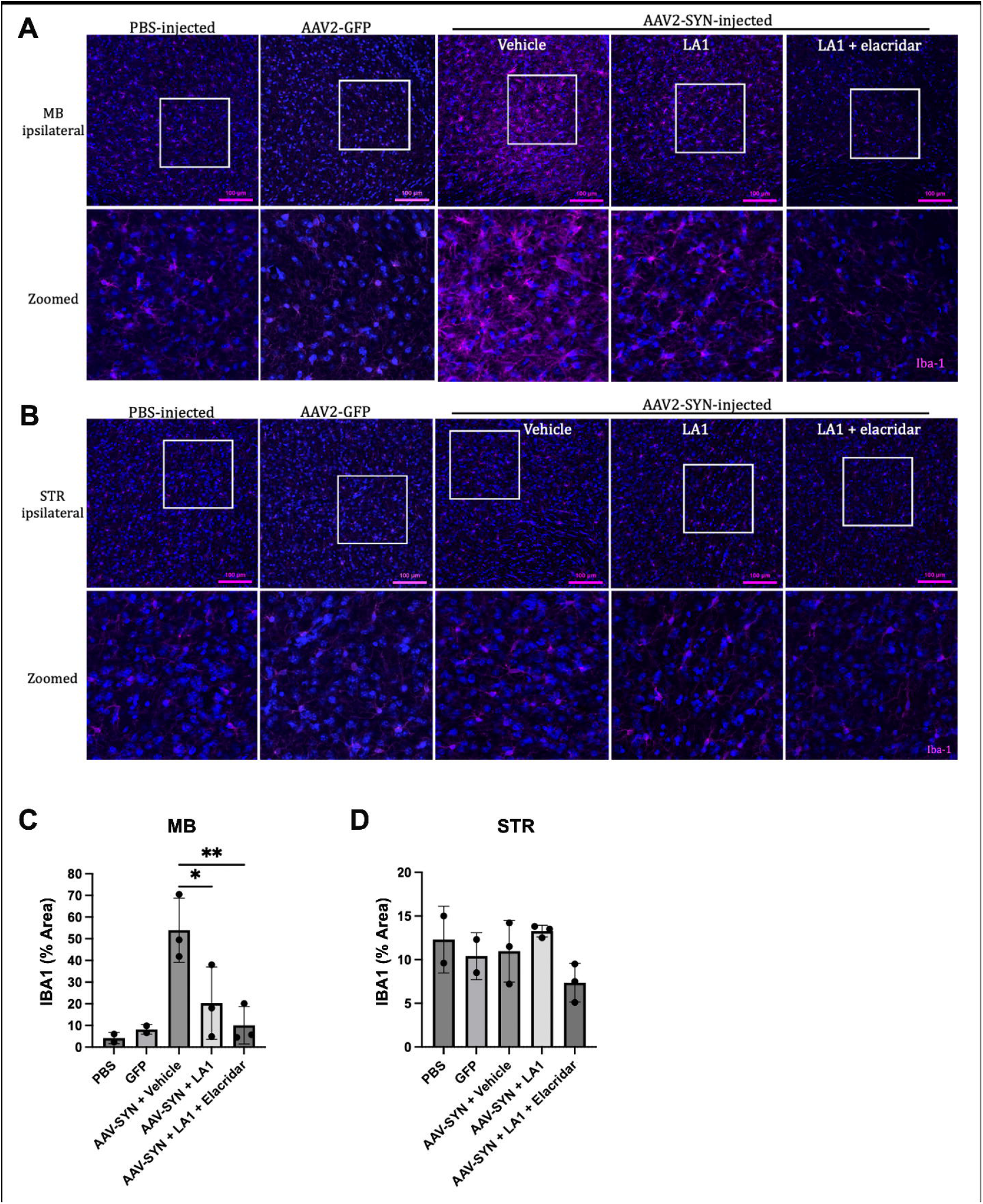
Iba-1 expression decreases in the midbrain after four weeks of treatment. Representative IF images of Iba-1 expression in the ipsilateral MB (A) and STR (B) of PBS, AAV-GFP, or AAV-SYN injected mice treated with vehicle, LA1, or LA1 with elacridar for four weeks. Images taken at 25X magnification (top) and digitally zoomed (bottom). Scale bar = 100μm. Subsequent quantification of Iba-1 immunoreactivity in the MB (C) and STR (D). Mean ± SEM is shown. PBS and AAV-GFP n = 2, AAV-SYN n = 3 per treatment group. *p<0.05, **p<0.01. MB = midbrain, STR = striatum.

A similar pattern was observed following eight weeks of treatment (Fig. 5). AAV-SYN mice receiving vehicle maintained elevated Iba-1 expression and activated microglial morphology in the midbrain (Fig. 5A, C). This response was significantly attenuated by LA1 treatment, and further diminished in mice treated with the combination of LA1 and elacridar, which restored Iba-1 expression and microglial morphology to levels comparable to PBS and AAV-GFP controls (Fig. 5A, C). As with the earlier time point, Iba-1 expression and microglial morphology in the striatum remained unchanged across all groups at eight weeks (Fig. 5B, D).

**FIGURE 5.**
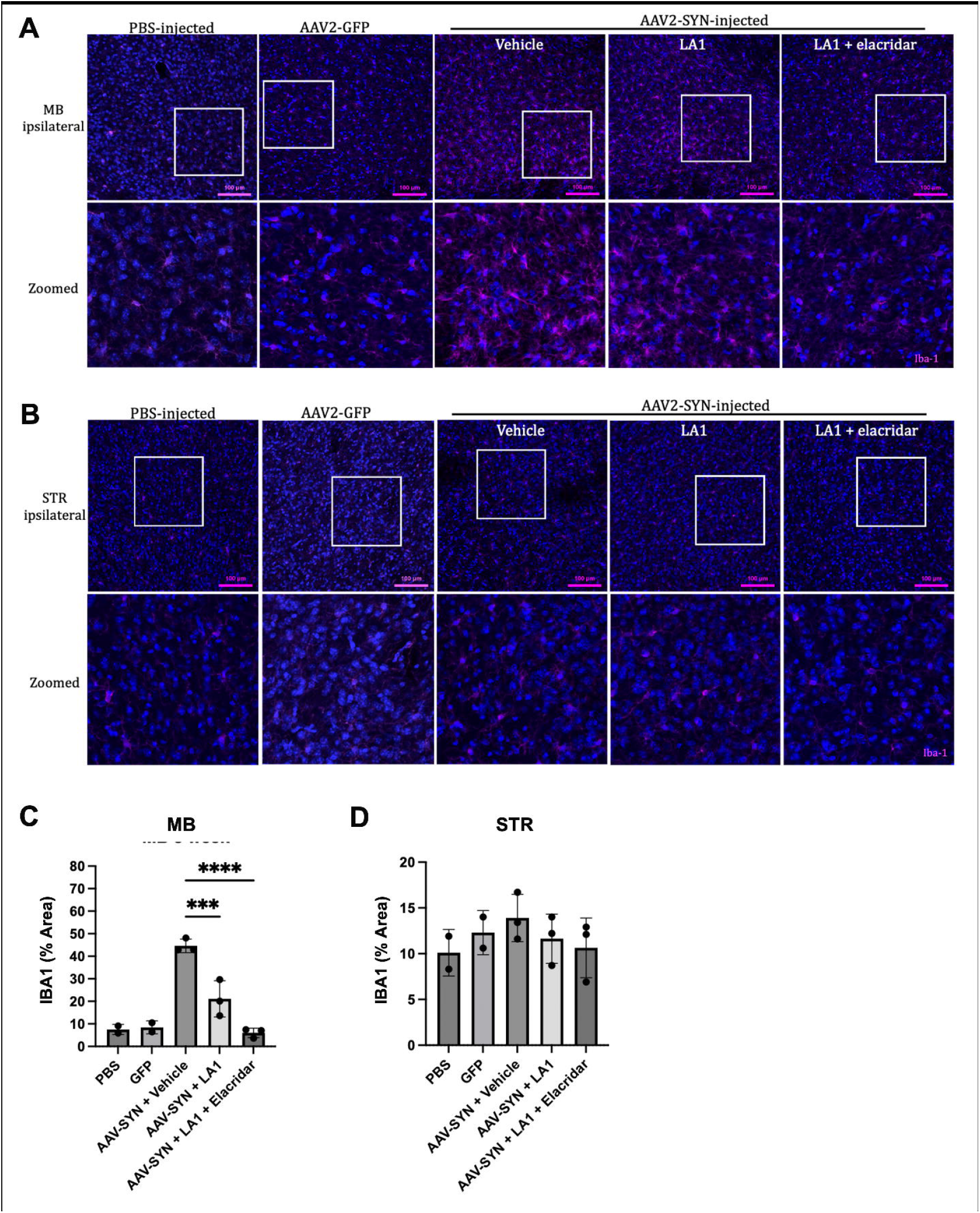
Iba-1 expression decreases in the midbrain after eight weeks of treatment. Representative IF images of Iba-1 expression in the ipsilateral MB (A) and STR (B) of PBS, AAV-GFP, or AAV-SYN injected mice treated with vehicle, LA1, or LA1 with elacridar for eight weeks. Images taken at 25X magnification (top) and digitally zoomed (bottom). Scale bar = 100μm. Subsequent quantification of Iba-1 immunoreactivity in the MB (C) and STR (D). Mean ± SEM is shown. AAV-GFP n = 2, PBS, AAV-SYN n = 3 per treatment group. ***p<0.0005, ****p<0.0001. MB = midbrain, STR = striatum.

## DISCUSSION

This study demonstrates that CD11b agonism via LA1 significantly mitigates α-synuclein–induced neuroinflammation in the AAV-SYN mouse model of Parkinson’s disease. Through a combination of pharmacokinetic optimization and targeted immunomodulation, we identify CD11b as a promising therapeutic target for modulating CNS inflammation in synucleinopathies. These findings support the therapeutic potential of CD11b agonists to shift innate immune responses and attenuate disease-associated pathology in PD.

Orally administered LA1 achieved measurable systemic exposure, however, CNS penetration of LA1 was limited, as evidenced by markedly lower brain concentrations relative to plasma. This low brain accumulation is consistent with active efflux via P-glycoprotein, a major efflux transporter at the blood-brain barrier known to restrict CNS entry of small molecules (22,23,24). Coadministration of elacridar, a P-gp inhibitor, significantly increased brain levels of LA1, achieving concentrations approaching those observed in plasma. These data suggest that LA1 is a substrate for P-gp and that inhibition of efflux transporters can enhance CNS bioavailability. Improved delivery to brain-resident immune cells in the presence of elacridar likely contributed to the enhanced anti-inflammatory effects observed in the combination treatment group. Such pharmacological strategies will be critical for translating CD11b-targeted immunotherapies to clinical use in neurodegenerative disease.

A hallmark of PD pathogenesis is the phosphorylation and accumulation of α-synuclein, particularly at serine 129 (pSer129), which is associated with Lewy pathology and neurodegeneration (25,26). In the AAV-SYN model, vehicle-treated mice exhibited robust nuclear accumulation of pSer129 in the midbrain, recapitulating features of PD-like pathology. Treatment with LA1 significantly reduced pSer129 expression and nuclear localization at both four and eight weeks, with an even greater reduction observed with elacridar coadministration. These findings suggest that CD11b agonism may reduce α-synuclein pathology either by dampening the proinflammatory milieu that promotes phosphorylation and aggregation, or by enhancing microglial clearance of misfolded protein. Given that neuroinflammation amplifies α-synuclein toxicity via cytokine signaling and oxidative stress (27), these results highlight the potential of immunomodulation to disrupt a key feed-forward loop in PD pathogenesis. Interestingly, pSer129 pathology was largely absent in the striatum despite viral α-synuclein overexpression, indicating regional differences in vulnerability that may be due to vector spread, neuronal subtype sensitivity, or intrinsic inflammatory tone (28). Nonetheless, the consistent reduction of pSer129 in the midbrain supports CD11b agonism as a strategy to mitigate α-synuclein-associated neurodegeneration.

CD11b, an integrin subunit expressed on microglia and peripheral myeloid cells, plays a critical role in regulating leukocyte adhesion, migration, and phagocytic function (14,29). Prior work has shown that CD11b agonists such as LA1 suppress inflammatory signaling, promote clearance of apoptotic cells, and shift immune cell phenotypes toward regulatory states (30,31). Consistent with these effects, we observed pronounced microglial activation in vehicle-treated AAV-SYN mice, characterized by bushy morphology and increased Iba-1 expression. LA1 treatment reversed these phenotypic changes, restoring a more quiescent microglial state, and combination treatment with elacridar further normalized microglial features to resemble those of surgical and vector controls. These changes were restricted to the midbrain, with no significant alterations in microglial activation observed in the striatum, again underscoring the region-specific nature of the inflammatory response in this model.

Together, these findings establish CD11b agonism as a viable strategy to reduce α-synuclein pathology and neuroinflammation in a preclinical model of Parkinson’s disease. The data also shows that targeting CD11b on circulating myeloid cells significantly reduces their brain infiltration and local inflammation, reducing synucleinopathy and might be sufficient to restore homeostasis in the diseased brain. These data lay the foundation for further investigation of CD11b-targeted agents in synucleinopathies and other neurodegenerative disorders characterized by maladaptive immune activation.

## EXPERIMENTAL PROCEDURES

### LA1 and Elacridar Pharmacokinetic studies

C57BL/6J (#000664 Jackson Laboratories) were oral gavaged with LA1 (30mg/kg; formulation: 3mg/mL solution – 6mg/mL water solution added with equal volume of 1% carboxymethylcellulose with 0.04% Tween 80 in water) and at 15min, 30min, 1h, 2h, 4h, and 8h post-treatment, mice were humanely euthanized, and plasma, brain, and kidneys were collected and analyzed via mass-spectrometry at The Herbert Wertheim UF Scripps Institute for Biomedical Innovation & Technology core at the University of Florida for pharmacokinetics and half-life studies. Data was collected using a mass spectrometer funded by NIH grant number 1 S10OD030332-01.

For pretreatment with the P-glycoprotein (P-gp) inhibitor, elacridar, mice were given 100mg/kg elacridar (formulation: 10mg/mL Elacridar in 10/20/13/57 DMSO/Tween80/EtOH/water) via intraperitoneal (IP) injection 30min prior to oral LA1 treatment and PK studies being performed as mentioned above.

### Mice

Male and female C57BL/6J (#000664 Jackson Laboratories) were used for these studies. Although sex was not analyzed as a biological variable, male and female mice were evenly distributed across experimental groups. For pooled midbrain and striatum samples, tissues from both sexes were combined to ensure sufficient cell yield. All mice were housed in clean, filter-top cages under a 12-hour light/dark cycle at an ambient temperature of 25°C, with *ad libitum* access to food and water. All animal research protocols were approved by the Institutional Animal Care and Use Committee at Rush University.

### AAV2-SYN and AAV2-GFP vectors

Neuronal overexpression of α-synuclein or green fluorescent protein (GFP) in the substantia nigra pars compacta (SNpc) was achieved through adeno-associated virus (AAV)-mediated transduction. The rAAV vectors, rAAV-1CBA-IRES2eGFP-WPRE (CIGW, AAV2-GFP) and rAAV-1CBA-aSYN-IRES2eGFP-WPRE (CISGW, AAV2-α-SYN), were constructed and purified as described in detail previously (33). The University of Iowa Viral Vector Core manufactured and purified AAV2-GFP and AAV2-SYN. Both AAV2-GFP and AAV2-SYN were stereotaxically injected into the SNpc of mice at a titer of 2.7 × 1012 vg/mL (33,34).

### Stereotaxic surgery

Surgical procedures followed established protocols (35). Mice were anesthetized using inhaled isoflurane and secured in a stereotaxic frame for intracranial injection. A total of 2 μL of AAV2-SYN or AAV2-GFP was administered into the SNpc at a rate of 0.5 μL/min using a Hamilton syringe. To ensure adequate diffusion, the syringe was left in place for an additional 3 minutes before slow retraction. Mice intended for immunohistochemistry analysis received unilateral injections to allow intra-animal comparisons. Those designated for flow cytometry received bilateral injections. The coordinates used for targeting, relative to bregma, were: AP −3.2 mm, ML ±1.2 mm, and DV −4.6 mm from the dura.

### LA1 and Elacridar treatments

One week post AAV2-SYN transduction, mice were treated orally twice daily with either vehicle (1% carboxymethylcellulose with 0.04% Tween 80 in water, BID), LA1 (30mg/kg BID; formulation: 3mg/mL solution – 6mg/mL water solution added with equal volume of vehicle: 1% carboxymethylcellulose with 0.04% Tween 80 in water), or LA1 + elacridar (30mg/kg LA1 with 100mg/kg elacridar BID; formulation same as LA1) for four or eight weeks.

### Mouse samples

At either four- or eight-weeks following treatment initiation, mice were deeply anesthetized and euthanized for tissue collection, including brain, blood, and spleen. Blood and spleen were collected prior to perfusion. Peripheral blood was collected in BD Vacutainer™ blood collection tubes with K2 EDTA (REF#367863, FisherScientific).

Unilaterally injected animals used for IHC/IF underwent transcardial perfusion with 0.01 M PBS followed by 4% paraformaldehyde (PFA). Whole brains were extracted and post-fixed overnight at 4 °C in 4% PFA and were then cryoprotected by sequential immersion in 15% and 30% sucrose solutions. Brains were frozen with dry ice and sectioned at 40 μm thickness using a freezing-stage microtome. Sections were stored at −20 °C in cryoprotectant solution containing 30% sucrose and 30% ethylene glycol in PBS.

### Immunohistochemistry

For fluorescent immunohistochemical analysis, free-floating brain sections were initially washed in dilution media (DM; 5 mM Tris–HCl, pH 7.6, 150 mM NaCl, 0.05% Triton X-100) followed by heat-induced antigen retrieval (HIAR) using sodium citrate buffer (10 mM sodium citrate, 0.05% Tween-20, pH 6.0).

Tissues were mounted on superfrost glass slides and dried before being covered with #1.5 glass coverslips using FluoroShield mounting medium (Sigma-Aldrich). Detailed protocol available at protocols.io (https://doi.org/10.17504/protocols.io.x54v92dq4l3e/v1).

For chromogenic detection using diaminobenzidine (DAB) staining floating brain sections were washed in dilution media (DM; 50 mM Tris-HCl, pH 7.4, 150 mM NaCl, 0.5% Triton X-100), then incubated for 1 hour at room temperature in a peroxidase quenching/blocking solution composed of 0.3% hydrogen peroxide, 0.1% sodium azide, 3% goat serum, 2% BSA, and 0.4% Triton X-100 in DM. Following washes in DM, sections were incubated overnight at 4 °C with either anti-PSER129 antibody (Abcam, EP1536Y, ab51253; RRID: AB_869973) at 1:50,000 in blocking buffer. The following day, sections were rinsed in DM and incubated for 1 hour at room temperature with a biotinylated anti-rabbit (Vector Laboratories, BA-1000; RRID: AB_2313606) secondary antibody diluted 1:200 in blocking buffer. After additional washes in DM, tissues were incubated for 75 minutes at room temperature with the VECTASTAIN Elite ABC reagent (Vector Laboratories, Cat# PK-6100, RRID: AB_2336819) to amplify signal. Following ABC incubation, sections were washed with DM and subsequently in sodium acetate buffer (0.2 M imidazole, 1.0 M sodium acetate, pH 7.2). Immunoreactivity was visualized using a nickel-enhanced 3,3′-diaminobenzidine (DAB)-imidazole reaction. Sections were then washed again in sodium acetate and PBS (50 mM Tris-HCl, pH 7.2, 158 mM NaCl), mounted onto glass slides, and counterstained with methyl green. Slides were dehydrated through graded ethanol, cleared in xylene, and coverslipped using Cytoseal 60 (Fisher Scientific). A comprehensive version of this immunohistochemistry protocol is available on protocols.io (https://doi.org/https://doi.org/10.17504/protocols.io.8epv5x3mdg1b/v1).

### Microscope Imaging and Quantification

Images were acquired using a Nikon A1R inverted confocal microscope using a 20X or 25X objective. Each tissue section was annotated within a standardized 2000 × 2000-pixel bounding box. Auto-exposure and auto-white balance corrections were applied to brightfield images using NIS-Elements software. Fluorescent images were exported and processed using Adobe photoshop and Illustrator. To quantify pSer129 and Iba-1 signals, an RGB-based color threshold was optimized to detect dark black or Cy5 pixels and recorded via the Macro function in NIS-Elements. This threshold was uniformly applied across all images, regardless of treatment group (PBS, AAV2-GFP, AAV2-SYN, or drug conditions: vehicle, LA1, or LA1 plus elacridar) or brain region (midbrain or striatum) (Figure S2). n = 3–6 animals were examined per treatment group per time point. The percentage area of thresholded signal was exported and visualized using GraphPad Prism. Statistical analysis was conducted using One-Way ANOVA with Dunnett’s multiple comparisons test, unless specified otherwise.

### Statistical analysis

All statistical analyses were performed using GraphPad Prism 10 software (v10.5.0). Data were analyzed using one-way ANOVA with Dunnett’s multiple comparison test. Graphs display the individual values and mean ± SE, with \**p*< 0.05, \*\**p*< 0.01, \*\*\**p*< 0.0005, \*\*\*\**p*< 0.0001.

## ACKNOWLEDGEMENTS

We thank members of the Gupta and the Killinger laboratories for helpful discussions and technical help.

## AUTHOR CONTRIBUTIONS

AMS designed and performed experiments with help from BN, VJ, JD, SC, and BK; AMS, BN, and VJ analyzed data; BK and VG designed and supervised the studies and AMS and VG co-wrote the paper. All authors reviewed and approved the final version of the manuscript.

## FOOTNOTES

This work was supported in part by grant MJFF-022480 from Michael J Fox Foundation and grants R01DK084195, R01DK136297, R01CA244938 from the National Institutes of Health (NIH) to VG and with resources from the Rush University Medical Center and the University of Texas Medical Branch (UTMB).

## REFERENCES

1. Tysnes, O. B., & Storstein, A. (2017). Epidemiology of Parkinson’s disease. Journal of Neural Transmission, 124(8), 901–905.

2. Spillantini, M. G., Schmidt, M. L., Lee, V. M., Trojanowski, J. Q., Jakes, R., & Goedert, M. (1997). α-Synuclein in Lewy bodies. Nature, 388(6645), 839–840.

3. Dickson, D. W. (2018). Neuropathology of Parkinson disease. Parkinsonism & Related Disorders, 46, S30–S33.

4. Poewe, W., Seppi, K., Tanner, C. M., et al. (2017). Parkinson disease. Nature Reviews Disease Primers, 3(1), 1–21.

5. Tansey, M. G., Wallings, R. L., Houser, M. C., Herrick, M. K., Keating, C. E., & Joers, V. (2022). Inflammation and immune dysfunction in Parkinson disease. Nature Reviews Immunology, 22, 657–673.

6. Hirsch, E. C., & Hunot, S. (2009). Neuroinflammation in Parkinson’s disease: A target for neuroprotection? The Lancet Neurology, 8(4), 382–397.

7. Subramaniam, S. R., & Federoff, H. J. (2017). Targeting microglial activation states as a therapeutic avenue in Parkinson’s disease. Frontiers in Aging Neuroscience, 9, 176.

8. Block, M. L., Zecca, L., & Hong, J. S. (2007). Microglia-mediated neurotoxicity: uncovering the molecular mechanisms. Nature Reviews Neuroscience, 8(1), 57–69.

9. Tang, Y., & Le, W. (2016). Differential roles of M1 and M2 microglia in neurodegenerative diseases. Molecular Neurobiology, 53(2), 1181–1194.

10. Tansey, M. G., & Romero-Ramos, M. (2019). Immune system responses in Parkinson’s disease: Early and dynamic. European Journal of Neuroscience, 49(3), 364–383.

11. Harms, A. S., Thome, A. D., Yan, Z., Schonhoff, A. M., Williams, G. P., Li, X., … & Tansey, M. G. (2018). Peripheral monocyte entry is required for α-synuclein induced inflammation and neurodegeneration in a model of Parkinson disease. Experimental Neurology, 300, 179–187.

12. Grozdanov, V., Bliederhaeuser, C., Ruf, W. P., Roth, V., Fundel-Clemens, K., Zondler, L., … & Danzer, K. M. (2014). Inflammatory dysregulation of blood monocytes in Parkinson’s disease patients. Acta Neuropathologica, 128, 651–663.

13. Kierdorf, K., & Prinz, M. (2017). Microglia in steady state. Journal of Clinical Investigation, 127(9), 3201–3209.

14. Ross, G. D., & Vetvicka, V. (1993). CR3 (CD11b, CD18): A phagocyte and NK cell membrane receptor with multiple ligand specificities and functions. Clinical and Experimental Immunology, 92(2), 181–184.

15. Ehirchiou, D., Xiong, Y., Xu, G., Chen, W., Shi, Y., & Zhang, L. (2007). CD11b facilitates the development of peripheral tolerance by suppressing Th17 differentiation. The Journal of Experimental Medicine, 204(7), 1519–1524.

16. Han, C., Jin, J., Xu, S., Liu, H., Li, N., & Cao, X. (2010). Integrin CD11b negatively regulates TLR-triggered inflammatory responses by activating Syk and promoting degradation of MyD88 and TRIF via Cbl-b. Nature Immunology, 11(8), 734–742.

17. Hemmati, S., Sadeghi, M. A., Yousefi-Manesh, H., Eslamiyeh, M., Vafaei, A., Foroutani, L., Donyadideh, G., Dehpour, A., & Rezaei, N. (2020). Protective effects of leukadherin1 in a rat model of targeted experimental autoimmune encephalomyelitis (EAE): possible role of P47phox and MDA downregulation. Journal of Inflammation Research, 411–420.

18. Yao, X., Dong, G., Zhu, Y., Yan, F., Zhang, H., Ma, Q., … & Si, C. (2019). Leukadherin-1-mediated activation of CD11b inhibits LPS-induced pro-inflammatory response in macrophages and protects mice against endotoxic shock by blocking LPS-TLR4 interaction. Frontiers in Immunology, 10, 215.

19. Faridi MH, Khan SQ, Zhao W, Lee HW, Altintas MM, Zhang K, et al. (2017). CD11b activation suppresses TLR-dependent inflammation and autoimmunity in systemic lupus erythematosus. Journal of Clinical Investigation, 127(4):1271–83.

20. Khan, S. Q., Khan, I., & Gupta, V. (2018). CD11b activity modulates pathogenesis of lupus nephritis. Frontiers in Medicine, 5, 52.

21. Geraghty, T., Rajagopalan, A., Aslam, R., Pohlman, A., Venkatesh, I., Zloza, A., Cimbaluk, D., DeNardo, D.G., & Gupta, V. (2020). Positive allosteric modulation of CD11b as a novel therapeutic strategy against lung cancer. Frontiers in Oncology, 10, 748.

22. Schinkel, A. H. (1999). P-glycoprotein, a gatekeeper in the blood–brain barrier. Advanced Drug Delivery Reviews, 36(2–3), 179–194.

23. Löscher, W., & Potschka, H. (2005). Blood-brain barrier active efflux transporters: ATP-binding cassette gene family. NeuroRx, 2(1), 86–98.

24. Miller, D. S., Bauer, B., & Hartz, A. M. (2008). Modulation of P-glycoprotein at the blood-brain barrier: opportunities to improve central nervous system pharmacotherapy. Pharmacological reviews, 60(2), 196–209.

25. Anderson, J. P., Walker, D.E., Goldstein, J.M., de Laat, R., Banducci, K., Caccavello, R.J., Barbour, R., Huang, J., Kling, K., Lee, M., Diep, L., Keim, P.S., Shen, X., Chataway, T., Schlossmacher, M.G., Seubert, P., Schenk, D., Sinha, S., Gai, W.P., Chilcote, T.J. (2006). Phosphorylation of Ser-129 is the dominant pathological modification of alpha-synuclein in familial and sporadic Lewy body disease. Journal of Biological Chemistry, 281(40), 29739–29752.

26. Fujiwara, H., Hasegawa, M., Dohmae, N., Kawashima, A., Masliah, E., Goldberg, M.S., Shen, J., Takio, K., Iwatsubo, T. (2002). alpha-Synuclein is phosphorylated in synucleinopathy lesions. Nature Cell Biology, 4(2), 160–164.

27. Harms, A. S., Delic, V., Thome, A. D., Bryant, N., Liu, Z., Chandra, S., Jurkuvenaite, A., West, A.B. (2017). α-Synuclein fibrils recruit peripheral immune cells via an endoplasmic reticulum stress pathway. Acta Neuropathologica Communications, 5(1), 85.

28. Ross, G. D. (2000). Regulation of the Adhesion versus Cytotoxic Functions of the Mac-1/CR3/α M β 2-lntegrin Glycoprotein. Critical Reviews™ in Immunology, 20(3).

29. Schittenhelm, L., Hilkens, C. M., & Morrison, V. L. (2017). β2 integrins as regulators of dendritic cell, monocyte, and macrophage function. Frontiers in Immunology, 8, 1866.

30. Schmid, M. C., Khan, S. Q., Kaneda, M. M., Pathria, P., Shepard, R., Louis, T. L., … & Varner, J. A. (2018). Integrin CD11b activation drives anti-tumor innate immunity. Nature communications, 9(1), 5379.

31. Li, Y., Ritzel, R. M., He, J., Liu, S., Zhang, L., & Wu, J. (2024). Ablation of the Integrin CD11b Mac 1 Limits Deleterious Responses to Traumatic Spinal Cord Injury and Improves Functional Recovery in Mice. Cells, 13(18), 1584.

