## Supplementary figures and images for "CD11b Activation Reduces Myeloid Brain Infiltration and Mitigates Synucleinopathy in a Model of Parkinson’s Disease"

### Shaw etal BiorXiv Aug2025 - supplemental figure 1.docx

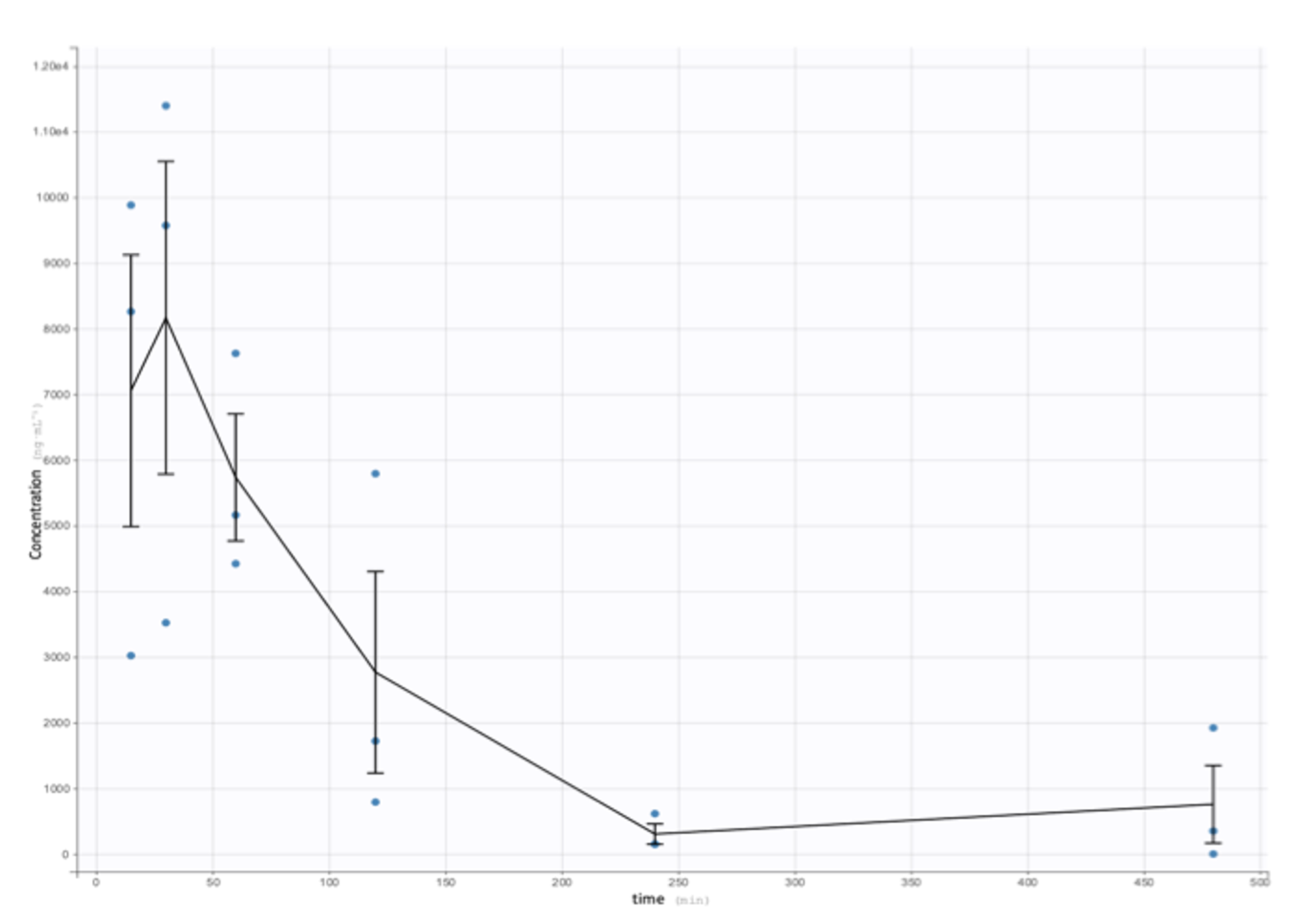

**FIGURE S1.** Pharmacokinetics of oral LA1. Mice received 30mg/kg LA-1 PO (oral). n=3/group/time-point.

### Shaw etal BiorXiv Aug2025 - supplemental table 2.docx

**TABLE S2.** LA1 concentration in plasma and brain after eight weeks of treatment.

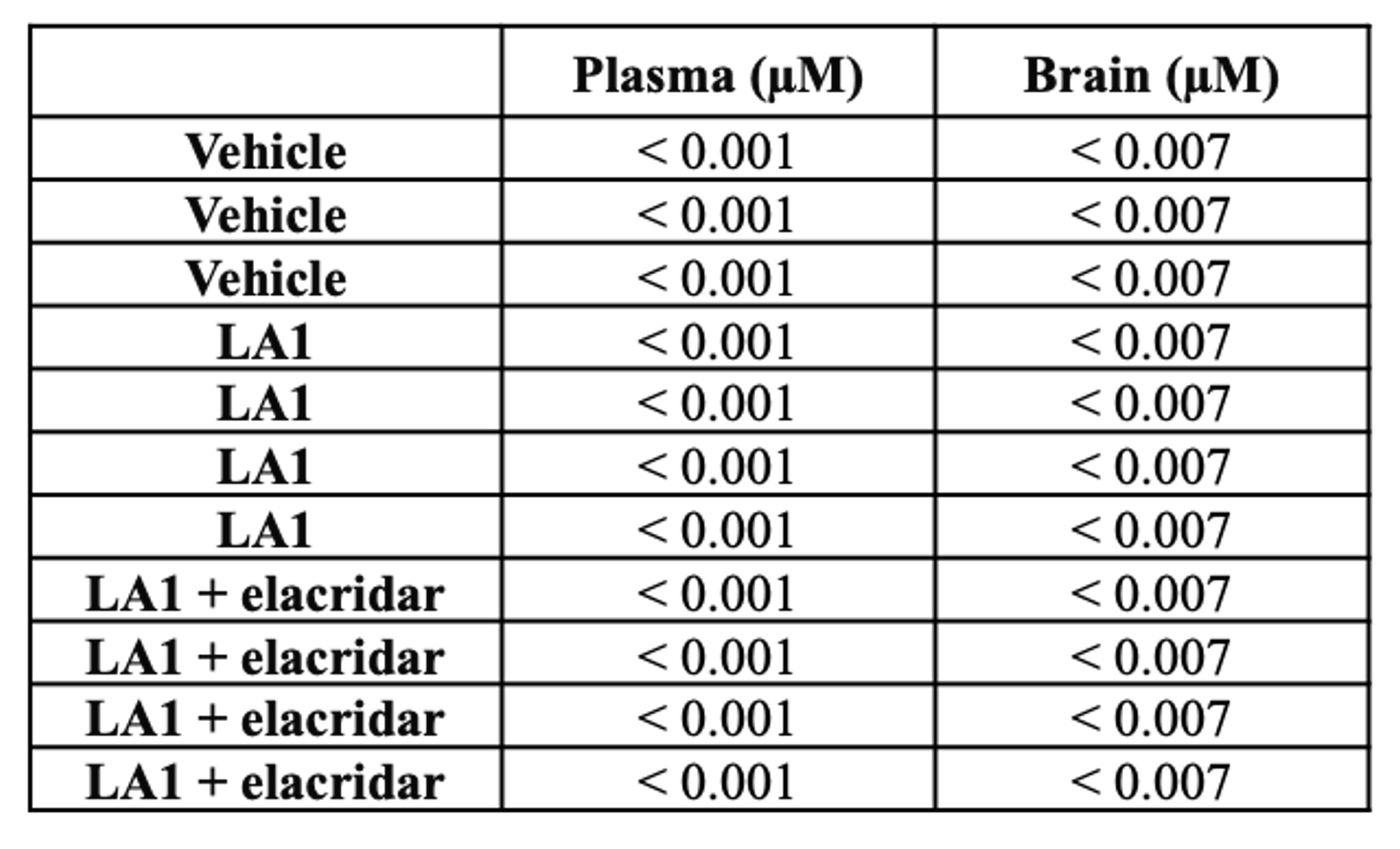
