## Supplemental Figure 2 for "CD11b Activation Reduces Myeloid Brain Infiltration and Mitigates Synucleinopathy in a Model of Parkinson’s Disease": Shaw etal BiorXiv Aug2025 - supplemental figure 2.docx

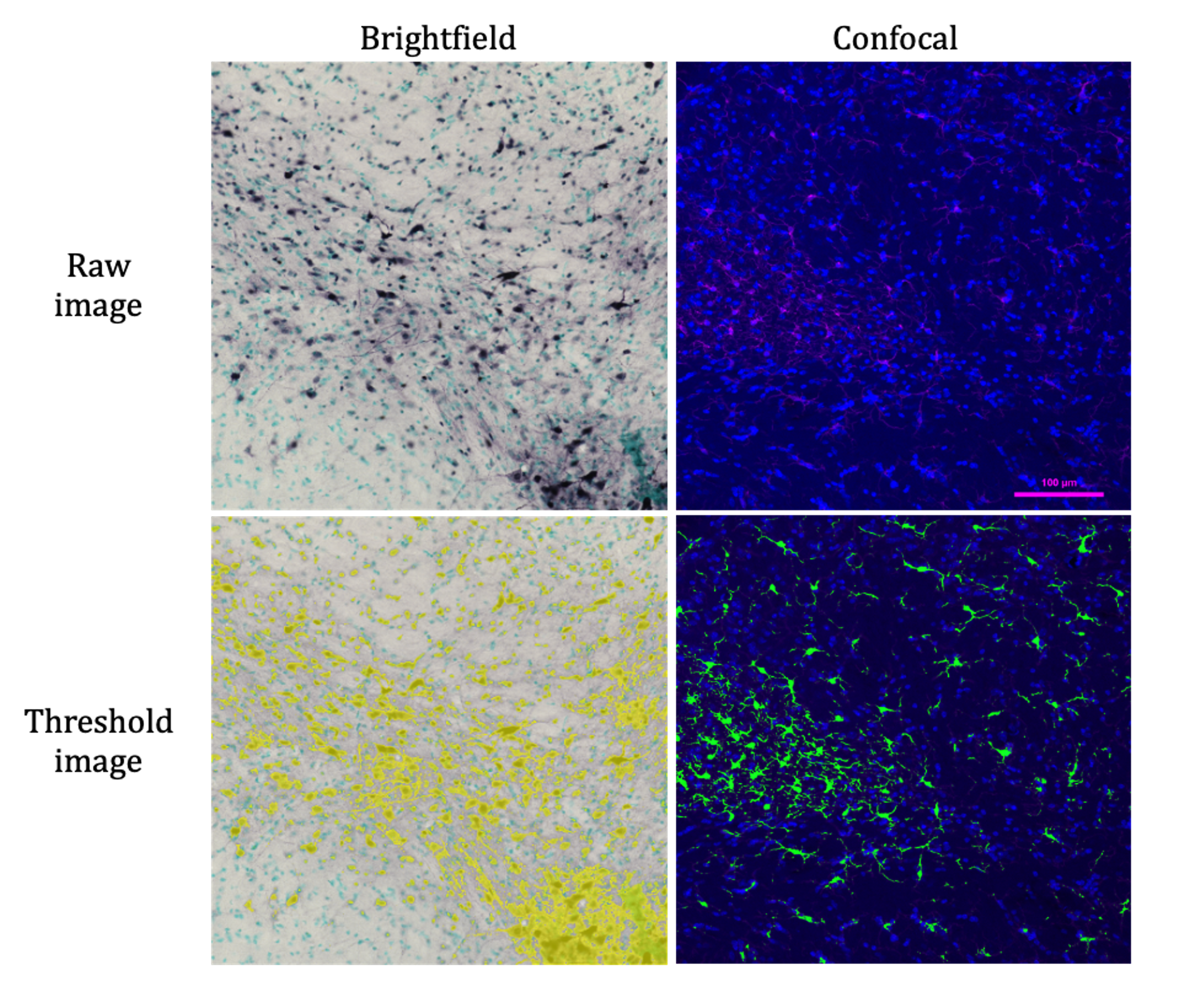

**FIGURE S2.** Thresholding examples of brightfield and confocal images. Left: pSer129 DAB IHC images. Right: Iba-1 IF images. Yellow and green indicates thresholded signals.
