## Supplemental Table 1 for "CD11b Activation Reduces Myeloid Brain Infiltration and Mitigates Synucleinopathy in a Model of Parkinson’s Disease": Shaw etal BiorXiv Aug2025 - supplemental table 1.docx

**TABLE S1.** Oral LA1 plasma pharmacokinetics. Mice received 30mg/kg LA-1 PO (oral). n=3/group/time-point. T_1/2_ = half-life, T_max_ = time to peak drug concentration, C_max_ = maximum concentration, AUC_last_ = area under curve to last measurable concentration, AUC_INF obs_ = area from time of dosing extrapolated to infinity, AUC_%Extrap_ = area under the curve extrapolated as a percentage of the total, Cl_obs = systemic clearance.

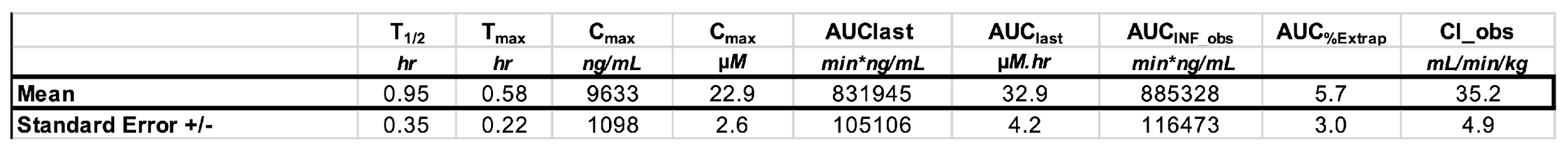
